# IFT20 regulates lymphatic endothelial cell-cell junctions via endocytic trafficking of VE-cadherin

**DOI:** 10.1101/2025.01.15.631989

**Authors:** Delayna Paulson, Ahana Majumder, Zachary Lehmann, Luke Knutson, Jacob Paulson, Shannon Lasey, Darci M. Fink

## Abstract

Intraflagellar transport (IFT) proteins are required for the assembly and function of primary cilia. They also regulate non-ciliary polarized vesicular traffic, such as T cell receptor recycling. We recently reported that lymphatic endothelial cells assemble primary cilia and express IFT proteins. Here, we report that IFT20 regulates vascular endothelial cadherin (VE-cadherin) localization at adherens junctions. IFT20 deletion caused discontinuous, button-like interendothelial junctions. This resulted in excessive lymphangiogenesis and impaired lymph drainage in mice. In vitro, VEGF-C treatment of IFT20 KD primary human dermal lymphatic endothelial cells caused accumulation of VE-cadherin in RAB5+ endosomes and enhanced and sustained VEGFR-3 signaling. Our findings are consistent with a model in which IFT20 promotes recycling of VE-cadherin to the adherens junction where it sequesters VEGFR-3 at the cell surface, thereby limiting pro-lymphangiogenic signaling. In the absence of IFT20, intercellular junctions are destabilized, pro-lymphangiogenic VEGFR-3 signaling is enhanced, and lymph transport is impaired by intracellular sequestration of VE-cadherin. This study elucidates the function of an IFT protein in lymphatic endothelial cells and provides mechanistic insight into the processes that regulate lymphatic endothelial cell-cell junctions and lymphangiogenic signaling.

## Introduction

Lymphatic endothelial cell-cell adhesion regulates lymphatic vessel permeability, lymph transport efficiency, and lymphangiogenesis. Together, these govern tissue fluid homeostasis. Lymphatic vessels can be subdivided into at least two classes with distinct junctional characteristics: capillaries and collecting vessels. Capillaries are specialized for uptake of fluid and cells and thus exhibit button-like junctions consisting of irregular patches of junctional proteins interspersed with flap-like openings(Baluk et al., 2007; Lynch et al., 2007; Pflicke and Sixt, 2009). Downstream, larger collecting vessels transport lymph to lymph nodes and back to the blood circulation. In collecting vessels, continuous, zipper-like intercellular junctions restrict permeability, and fluid transport is supported by rhythmic contractions of lymphatic muscle cells that propel lymph unidirectionally through lymphatic valves(Weid and Zawieja, 2004). The button- vs. zipper-like distribution of junctional constituents such as vascular endothelial cadherin (VE-cadherin) and zonula occludens proteins is dynamically regulated during vascular maturation and is responsive to tissue microenvironment cues such as increased fluid shear stress or inflammatory signals(Kelley et al., 2011; Yao et al., 2012; Zheng et al., 2014). Disruption of VE-cadherin continuity increases permeability of lymphatic vessels(Jannaway and Scallan, 2021) and promotes lymphangiogenic vascular endothelial growth factor C (VEGF-C) signaling(Sung et al., 2022). During acute lymphangiogenesis, VEGF-C stimulates quiescent lymphatic endothelial cells (LECs) in pre-existing vessels to sprout, migrate, and proliferate to expand the lymphatic plexus(Kelley et al., 2013; Deng et al., 2015). In this study, we identified an intraflagellar transport protein, IFT20, as a novel regulator of lymphatic cell-cell junctions, with consequences for lymphangiogenesis and edema.

Intraflagellar transport (IFT) proteins form complexes with motor proteins to carry cargo along the cytoskeletal axoneme of cilia(Kozminski et al., 1993, 1995; Hesketh et al., 2022; Lacey et al., 2023, 2024). The primary cilium is a non-motile signaling organelle that extends from the plasma membrane and houses specific membranous and cilioplasmic receptors and effector proteins within its unique subcellular microenvironment. Deletion of specific IFT proteins impairs primary cilia assembly and function and is a widely used model to study cell type-specific primary cilia signaling and mechanosensing(Pazour et al., 2000; Jonassen et al., 2008; Wang et al., 2021; Bakey et al., 2023). We recently reported that LECs exhibit primary cilia *in vitro* and *in vivo*(Paulson et al., 2021). Deletion of IFT20 prevented assembly of primary cilia and caused defects in lymphatic vessel patterning during mouse embryonic development and corneal inflammation, demonstrating the importance of this system in LEC biology. Mouse embryos with global IFT20 deletion exhibited enlarged dermal lymphatics with hyperbranching and severe edema. In adult mice, lymphatic-specific KO of IFT20 exacerbated suture-induced corneal lymphangiogenesis. These data suggested that IFT20 restricts lymphatic vessel growth and promotes lymph transport.

IFT20 is a 15 kDa protein that associates with other IFT proteins in IFT complex B to promote anterograde transport of cargo proteins up the ciliary axoneme(Lacey et al., 2023). IFT20 also localizes to the Golgi apparatus, where it regulates polarized vesicular trafficking of ciliary proteins to the primary cilium(Follit et al., 2006, 2008; Monis et al., 2017). Recent reports have demonstrated many additional primary cilium-independent trafficking functions of IFT20. Notably, IFT20 is required for recycling of the T cell receptor at the immune synapse(Finetti et al., 2009, 2014, 2015; Vivar et al., 2016) and turnover of integrins at focal adhesions(Su et al., 2020), two processes that, like transport to the cilium, rely on polarized vesicular traffic to specific plasma membrane regions. Here, we tested the hypothesis that IFT20 regulates lymphatic vessel permeability and lymphangiogenesis by directing polarized recycling of VE-cadherin to the adherens junction. We show that depletion of IFT20 causes accumulation of VE-cadherin in RAB5+ endosomes and concomitant breakdown in intercellular junctions. This prevents efficient lymph drainage and intensifies lymphangiogenesis by increasing VEGF-C signaling. This study identifies a novel mechanism of intercellular junction and growth factor signaling regulation in LECs that requires a nonciliary trafficking role of IFT20.

## Results

### Lymphatic-specific IFT20 KO increases suture-induced lymphangiogenesis and causes discontinuous VE-cadherin localization at intercellular junctions

To interrogate the effects of loss of IFT20 on inflammation-induced lymphangiogenesis *in vivo*, we used LYVE1Cre(Pham et al., 2010) to delete IFT20 from the lymphatic vasculature. We employed a suture-induced model of corneal inflammation to induce the growth of new lymphatic and blood vessels in a healthy avascular cornea(Kelley et al., 2013). Four sutures were placed in the cornea of lymphatic-specific IFT20 KO mice (bearing one copy of LYVE1Cre and two floxed alleles of *Ift20*) and littermate control mice (negative for LYVE1Cre and bearing one floxed allele of *Ift20*) (**Figure 1A**). After 12 days, suture-induced vascularization was analyzed by immunofluorescence microscopy. Lymphatic vessels were identified by expression of LYVE-1 and CD31. Blood vessels expressed CD31 and were negative for LYVE-1. In lymphatic-specific IFT20 KO mice, excessive lymphangiogenesis and hemangiogenesis (due to slight leakiness of LYVE1Cre) was observed relative to littermate control (**Figure 1B, C, D**), similar to results we previously reported in a 7-day inflammation model(Paulson et al., 2021). These data suggest that IFT20 regulates lymphangiogenesis during corneal inflammation, a process that relies on VEGF-C signaling(Kelley et al., 2013).

**Figure 1.**
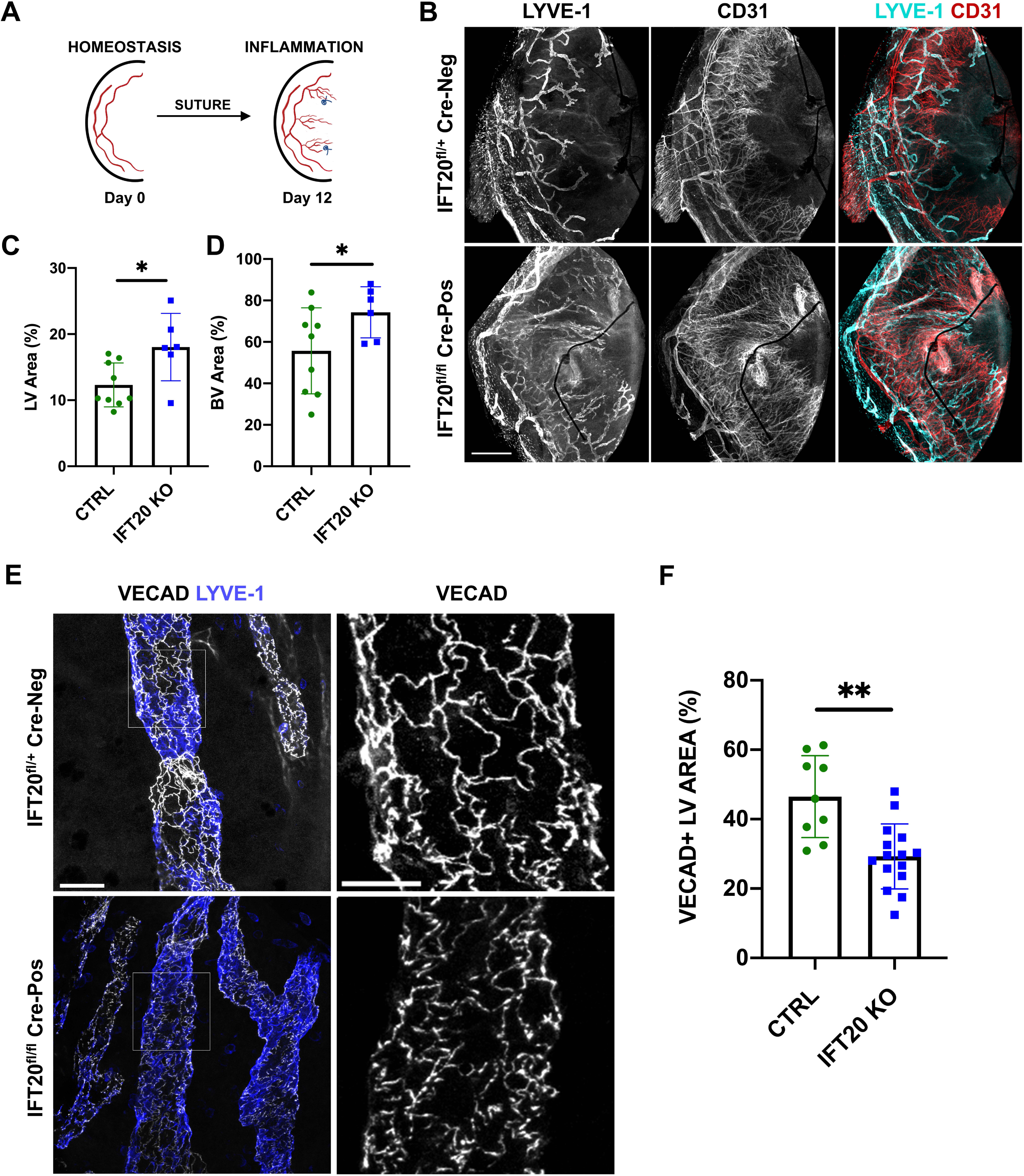
Lymphatic-specific IFT20 KO increases suture-induced lymphangiogenesis and causes discontinuous VE-cadherin localization at intercellular junctions. **(A)** Four sutures were placed in the healthy adult mouse cornea on day 0 to induce inflammation. Corneas were harvested on day 12 for immunofluorescence analysis. **(B)** Whole mount hemisected mouse corneas immunostained for LYVE-1 (*cyan*, lymphatic surface marker) and CD31 (*red,* pan-endothelial marker). LEC-specific IFT20 KO mice expressed LYVE1Cre and carried two floxed alleles of *Ift20*. Littermate controls were negative for Cre and carried one floxed allele of *Ift20*. Corneas were imaged using laser scanning confocal microscopy and micrographs shown are maximum intensity projections (MIPs). Scale bar = 500 μm. **(C)** Quantification of lymphatic vessel area as a percentage of total corneal area from *B*. **(D)** Quantification of blood vessel area as a percentage of total corneal area from *B*. **(E)** Whole mount immunofluorescence staining for VE-cadherin (*white,* VECAD) and LYVE-1 (*blue*) in lymphatic specific-IFT20 KO mice and littermate controls as in *B*. Corneas were imaged using laser scanning confocal microscopy. Micrographs shown are MIPs from z-stacks encompassing volumes that contain LYVE-1+ lymphatic vessels. Scale bar *left* = 50 μm. Scale bar *right* = 25 μm. **(F)** Quantification of VECAD+ lymphatic vessel (LV) area as a percentage of total LV area. **(C, D, F)** Each data point represents one field of view. For each quantification, two to four FOVs were quantified from each of four IFT20 KO mice and three negative control littermates. *p<0.05, **p<0.005.

Given that interendothelial junctions dynamically remodel in response to inflammation(Kelley et al., 2011; Yao et al., 2012; Zheng et al., 2014), we predicted that changes in cell-cell junctions might accompany the excessive lymphangiogenesis phenotype observed in IFT20 KO mice. VEGF-C/VEGFR-3 signaling is sufficient to drive corneal lymphangiogenesis(Cursiefen et al., 2004; Kelley et al., 2013). VEGFR-3 activation induces Src-mediated phosphorylation and internalization of VE-cadherin(Sung et al., 2022), a critical homotypic adhesion molecule at the endothelial adherens junction. Internalization of VE-cadherin disrupts endothelial cell-cell junctions, enhancing migratory capacity of individual cells and increasing vascular permeability. Lymphatic vessels at the periphery of the cornea respond to inflammatory stimuli by sprouting and proliferation to initiate expansion of the corneal lymphatic plexus(Cursiefen et al., 2003). These peripheral vessels exhibit patchy LYVE-1 expression, consistent with pre-collecting lymphatic vessels(Mäkinen et al., 2005). Immunofluorescence microscopy at the late stages of suture-induced lymphangiogenesis revealed that VE-cadherin displayed a punctate, button-like arrangement on LYVE-1+ endothelial cells (**Figure 1E**). In contrast, littermate controls had uniform VE-cadherin at the perimeter of LECs consistent with zipper junctions typically observed in pre-collector and collecting lymphatic vessels. Quantification supported this observation, with the area of VE-cadherin covering LYVE-1+ lymphatic vessel area reduced in IFT20 KO mice (**Figure 1F**). These results demonstrate that IFT20 is an important regulator of VE-cadherin localization at the adherens junction during VEGF-C-dependent corneal lymphangiogenesis and predict that IFT20 supports robust endothelial junctions.

### Lymph drainage is impaired in IFT20 KO skin

Disrupted VE-cadherin organization on IFT20 KO lymphatic vessels led us to investigate lymphatic function. We previously reported that deletion of IFT20 early in gestation caused pronounced embryonic edema(Paulson et al., 2021) that was very similar to edema caused by VE-cadherin deletion(Hägerling et al., 2018; Harris et al., 2022). Lymphatic vessels in the IFT20 KO or VE-cadherin KO embryonic dorsal skin exhibited increased and variable lumen size and excessive branching, suggesting that impaired lymphatic organization and function contributed to the fluid homeostasis defect. Here, immunofluorescence staining for LYVE-1 in the ear skin revealed similar patterning defects in adult IFT20 KO lymphatic vessels (**Figure 2A**), that have also been described in VE-cadherin KO mice(Hägerling et al., 2018). We hypothesized that lymphatic drainage function would be impaired in adult IFT20 KO mice. Injection of a tracer into the adult mouse ear dermis is a widely used model to track lymph transport by intravital imaging (**Figure 2B**). We injected TRITC-labeled dextran (MW 40kD) into the ear skin and observed fluorescent lymph uptake and transport by intravital microscopy over 20 min (**Figures 2C-E**). In littermate controls, fluorescent lymph entered lymphatic capillaries and was efficiently transported in the direction of the ear base through larger collecting vessels punctuated by lymphatic valves. However, lymph transport was impaired in IFT20 KO mice. IFT20 KO skin exhibited defects in lymphatic capillary organization as fluorescent lymph collected in irregularly shaped pockets of distended lumen rather than draining away (**Figure 2D**). Retrograde flow of lymph was observed in KOs, with sections of upstream lymphatic capillaries filling during and after injection, possibly indicative of a defect in lymphatic valve patency (**Figures 2C, E**). Strikingly, diffuse fluorescent dextran signal could be visualized in interstitial areas outside KO lymphatic vessels, including at locations distant from the injection site. This widespread extravascular signal was not observed in littermate control mice in which fluorescent dextran was localized to the draining lymphatic vessel network. These data demonstrate that IFT20 KO lymphatic vessels do not efficiently transport lymph due to pooling of fluid in abnormally-shaped capillary structures, retrograde flow, and leakage into the interstitium.

**Figure 2.**
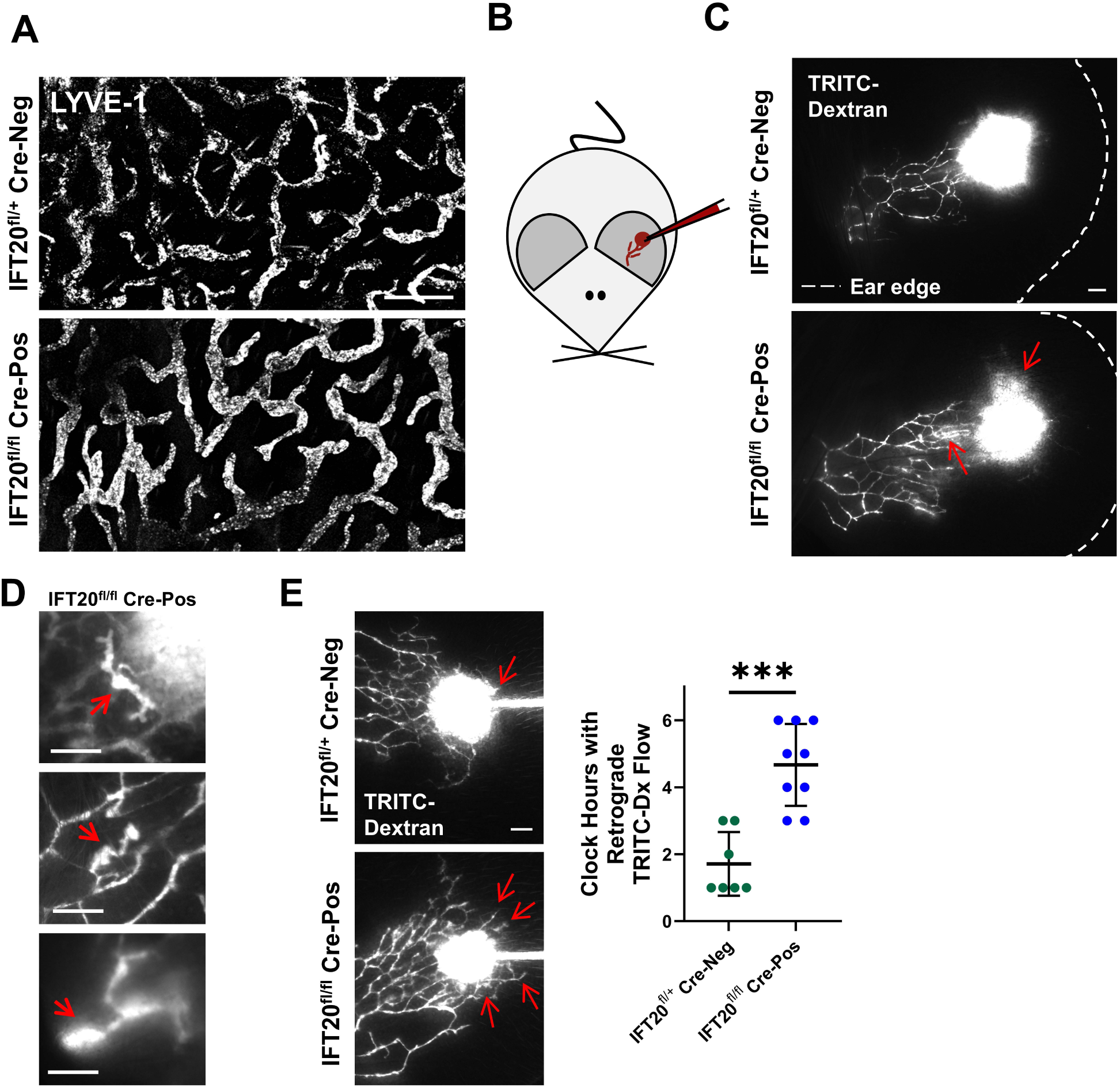
Lymphatic vessel drainage function is impaired in IFT20 KO skin. **(A)** Immunofluorescence micrographs of LYVE-1+ lymphatic vessel morphology in mouse ear skin. LEC-specific IFT20 KO mice expressed LYVE1Cre and carried two floxed alleles of *Ift20*. Littermate controls were negative for Cre and carried one floxed allele of *Ift20*. Scale bar = 100 μm. **(B)** To assess the ability of lymphatic vessels to transport lymph, 0.5 μL of TRITC-conjugated dextran (MW 40 kDa) was injected into mouse ear dermis using a stereotactic injector fitted with a pulled glass needle. Fluorescent lymph uptake and transport by lymphatic vessels was assessed by intravital stereofluorescence microscopy. **(C-E)** Intravital microscopy tracking fluorescent dextran drainage through ear dermis lymphatic vessels in lymphatic-specific IFT20 KO mice and Cre-negative littermate controls. *C* shows clean transport of TRITC-dextran toward the base of the ear in control mice, while IFT20 KO mice displayed retrograde flow (*red arrows*) and extravascular fluorescent signal in the interstitium. Images shown are from 20 min after injection. **(D)** *Red arrows* indicate accumulation of TRITC-dextran in abnormally-shaped lymphatic capillary structures. **(E)** *Red arrows* indicate retrograde flow, which is significantly more pronounced in IFT20 KOs. Quantification shows the number of clock hours (from 12 o’clock to 6 o’clock) per injection displaying retrograde flow. Quantification includes data from both 0.5 and 4 μL injections from 3 experiments in IFT20 KO and control mice. p = 0.0001. Scale bars = 100 μm.

### Depletion of IFT20 impairs intercellular junction formation and increases transwell migration of lymphatic endothelial cells

After establishing the *in vivo* relevance of IFT20 for lymphatic vessel patterning and function, we used a reductionist *in vitro* model to identify the IFT20-dependent cellular mechanisms by which LECs regulate VE-cadherin organization. We used CRISPR/Cas9 to knockout IFT20 in an immortalized mouse lymphatic endothelial cell line (mLEC) (Ando et al., 2005). Deletion of IFT20 was confirmed by immunofluorescence staining (**Figure 3A**). As we previously reported *in vivo*(Paulson et al., 2021), deletion of IFT20 significantly reduced the incidence of ARL13B+ primary cilia on 24 h serum-starved mLECs (**Figure 3A, B**). This indicates that IFT20 is required for primary cilia assembly on LECs and agrees with previous studies in which other cell types also failed to assemble primary cilia in the absence of IFT20(Follit et al., 2006; Jonassen et al., 2008). Using this model, we first sought to understand how loss of IFT20 would affect intercellular junctions between mLECs *in vitro* (**Figure 3C**). We labeled F-actin with phalloidin in confluent cultures of parental and IFT20 KO mLECs and found increased stress fiber formation and gaps between KO cells, while parental cells adhered tightly to one another. Immunofluorescence staining of zonula occludens-1 (ZO-1), a component of tight junctions, further demonstrated the dysregulation of intercellular junctions in IFT20 KO cells. The perimeter of parental cells showed linear continuous ZO-1 staining, while cell-cell contacts between KO cells were less clearly defined, and ZO-1 was localized in punctate structures at the cellular perimeter. The antibodies we used were unable to label VE-cadherin in these cells. Deletion of VE-cadherin itself has been reported to either upregulate, downregulate, or have no effect on expression and organization of other junctional proteins depending on the context(Taddei et al., 2008; Frye et al., 2015; Hägerling et al., 2018; Yang et al., 2019), so it is interesting but unsurprising that other junctional proteins are affected.

**Figure 3.**
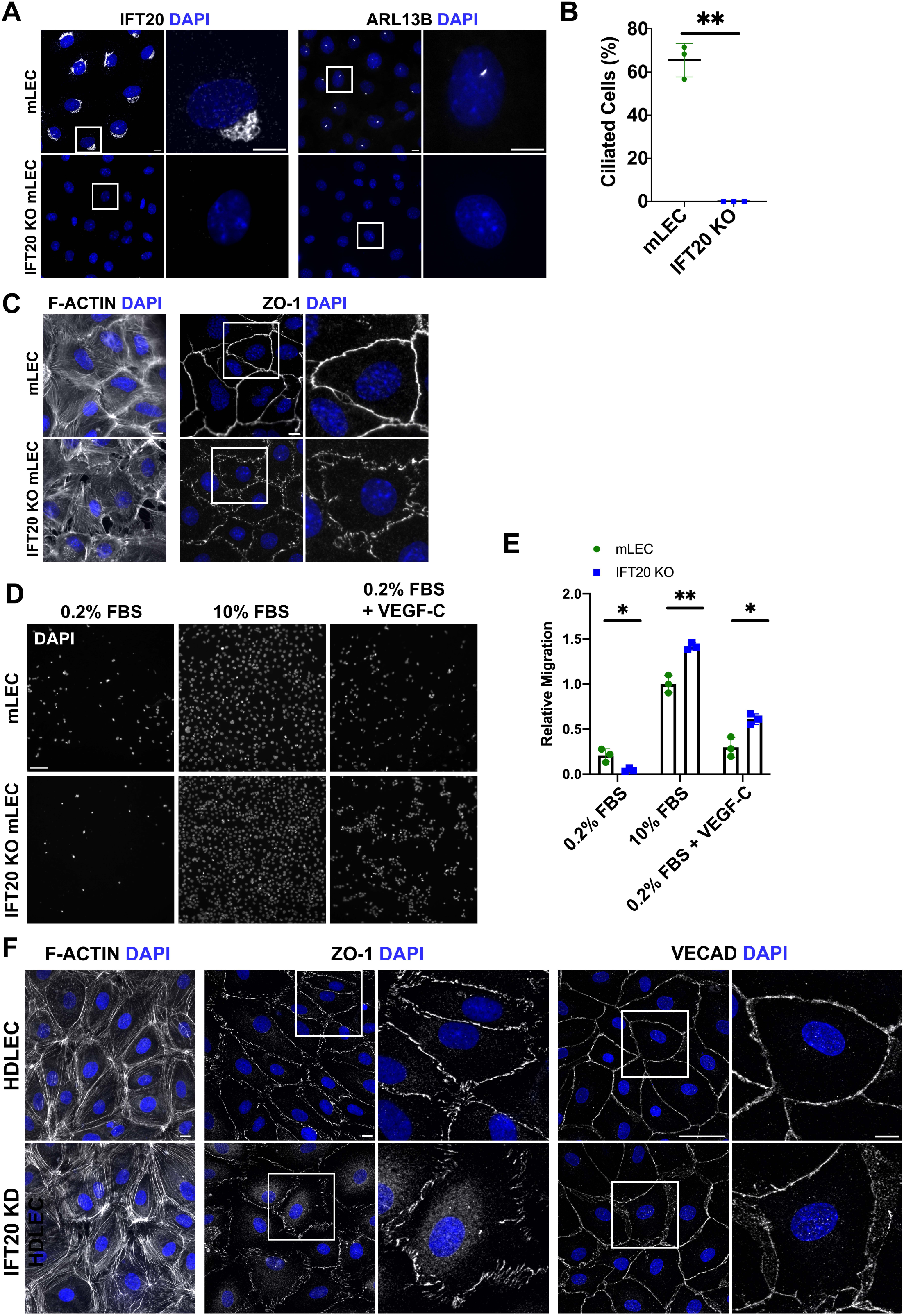
Depletion of IFT20 impairs intercellular junction formation and increases transwell migration of LECs. **(A)** *Ift20* was knocked out of immortalized mouse lymphatic endothelial cells (mLEC) using CRISPR/Cas9. Immunofluorescence micrographs confirm loss of IFT20 (*left panel*) and abrogation of primary cilia assembly in IFT20 KO cells after 24 h serum starvation in media containing 0.2% FBS (*right panel,* quantified in *B*). DAPI (*blue*). Scale bars = 10 μm. **(B)** Quantification of primary cilia incidence in parental and IFT20 KO mLECs from *A*. Data points represent each of three independent experiments, each quantifying 100+ cells. **(C)** Epifluorescence micrographs of parental and IFT20 KO mLECs with phalloidin-labeled F-actin (*left panel*) or ZO-1 immunofluorescence staining (*right panel*). DAPI (*blue*). Scale bars = 10 μm. **(D)** Chemotaxis potential of parental and IFT20 KO mLECs was assessed by transwell migration assay. Cells were seeded in the upper chamber in starving media containing 0.2% FBS. Lower wells were filled with media containing either 0.2% FBS, 10% FBS, or 500 ng/mL VEGF-C in 0.2% FBS. After 24 h migration, membranes were fixed, mounted in DAPI mounting media, and imaged via epifluorescence microscopy. Scale bar = 200 μm. **(E)** Quantification of transwell migration from *D*. Migration is graphed relative to 10% FBS-stimulated mLEC migration. Each data point represents an average from 5 FOVs per membrane. Quantification is representative of three independent experiments in which three membranes were quantified per experimental condition. **(F)** *Ift20* was targeted by siRNA in primary human dermal lymphatic endothelial cells (HDLECs). F-actin was labelled with phalloidin, and ZO-1 and VE-cadherin were labeled by immunofluorescence. Micrographs are MIPs from laser scanning confocal microscopy. DAPI (*blue*). Scale bars F-actin and ZO-1 = 10 μm. Scale bar VECAD *left panel* = 50 μm, *right panel* = 10 μm. *p<0.05, **p<0.005.

After establishing that this mouse IFT20 KO cell line displayed similar cell-cell adhesion defects to those observed *in vivo*, we next sought to determine how this would impact LEC migration. As discussed above, VEGF-C is the primary growth factor that drives lymphangiogenesis by signaling through its receptors VEGFR-2/3 on LECs (Deng et al., 2015; Monaghan et al., 2020). To determine whether loss of IFT20 affects this signaling axis, we performed transwell migration assays in the presence of VEGF-C as a chemoattractant (**Figure 3D, E**). As expected, parental mLECs were strongly attracted to media containing either positive control 10% FBS or 500 ng/mL VEGF-C + 0.2% FBS vs. 0.2% FBS media alone. IFT20 KO LECs migrated more than parental cells in response to both 10% FBS and VEGF-C, respectively. KO of VE-cadherin in primary mouse LECs similarly facilitated increased migration(Hägerling et al., 2018). This suggests that IFT20 negatively regulates LEC migration, and that the increased migration in IFT20 KO cells may be due to decreased intercellular adhesion between LECs.

Primary human dermal lymphatic endothelial cells (HDLECs) are the most physiologically relevant *in vitro* culture system currently available for lymphatic research. For the remainder of the studies reported here, we used commercially available adult HDLECs. These cells consist of a mixture of capillary and collector HDLECs that are isolated from the skin of healthy donors. We used siRNA to deplete IFT20 from these cells (please see confirmation of KD by immunofluorescence in Figure 4A and western blot in Figure 7A) and then used immunofluorescence to characterize the state of intercellular adhesion in control and IFT20 KD HDLECs (**Figure 3F**). In IFT20 KD HDLECs, F-actin labeling showed increased stress fiber formation and gaps between cells, ZO-1 was organized in patches with variable orientation relative to cellular perimeter, and VE-cadherin was dispersed and punctate. These results demonstrate that the impaired intercellular adhesion that we observed *in vivo* and *in vitro* with mouse LECs is also a feature of human IFT20 KD LECs. Taken together, these data suggest that IFT20 regulates the continuity of intercellular adherens and tight junctions across mammalian species, and that VEGF-C stimulation is sufficient to stimulate hypermigration in IFT20 KO cells.

**Figure 4.**
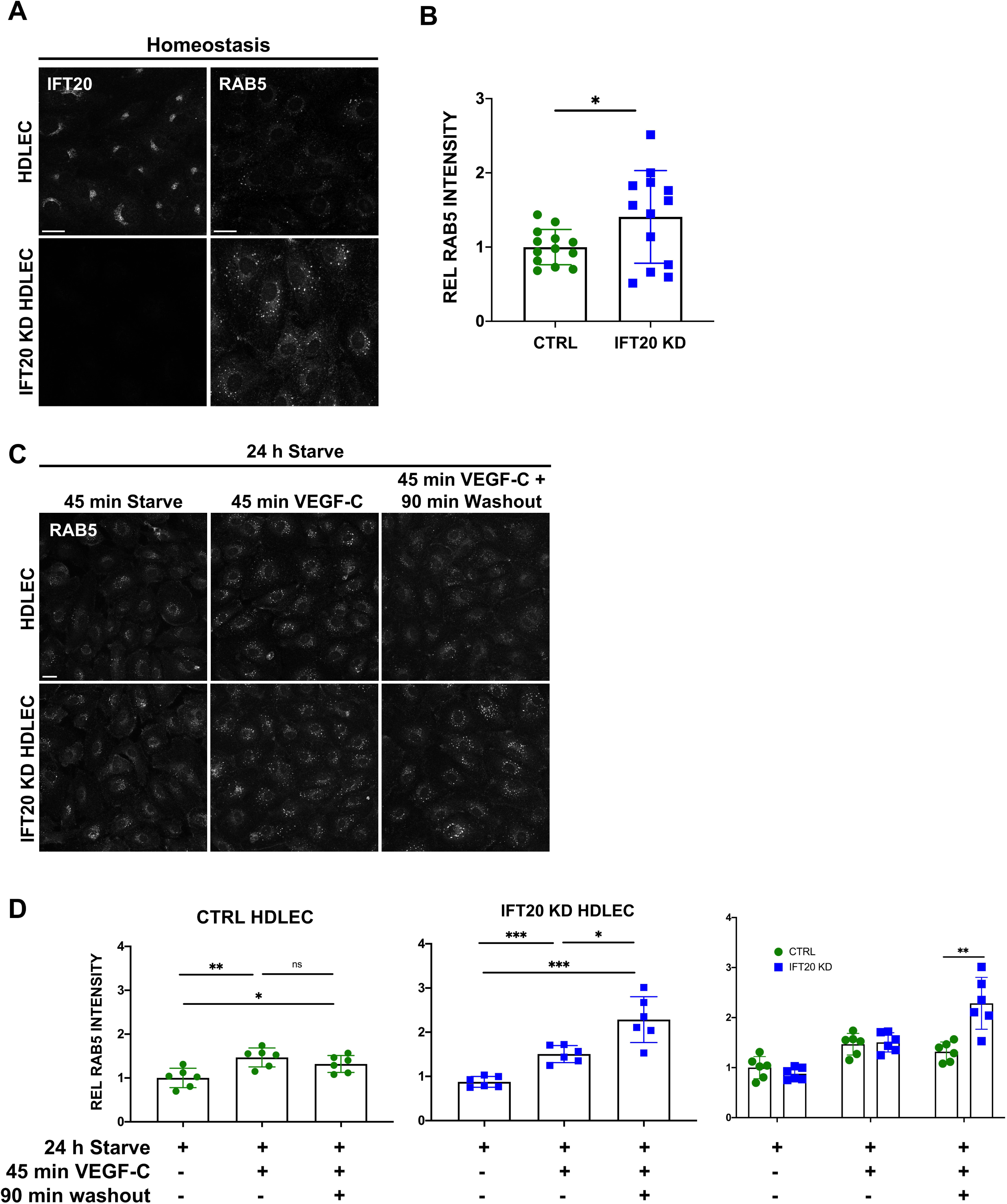
RAB5+ endosomes accumulate in IFT20 KD human dermal LECs. **(A)** Control and IFT20 KD HDLECs in homeostasis were immunostained for IFT20 and RAB5. Micrographs are MIPs from laser scanning confocal microscopy. Scale bars = 25 μm. **(B)** Quantification of integrated intensity from RAB5+ area in control and IFT20 KD HDLECs. Intensities are graphed relative to average control HDLEC RAB5 intensity. Quantification is from one of three biological replicates and is representative of 300+ control and 300+ IFT20 KD HDLECs. **(C)** Control and IFT20 KD HDLECs were serum starved in media containing 1/5 EGM-MV2 and 4/5 EBM-2 (hereafter, starving media) for 24 h. Cells were then either: placed in fresh starving media for 45 min, then fixed; placed in fresh starving media + 2 μg/mL VEGF-C for 45 min, then fixed; or placed in fresh starving media + 2 μg/mL VEGF-C for 45 min, then placed in EBM-2 basal media for a 90 min washout, then fixed. RAB5 was detected in fixed cells by immunofluorescence. Micrographs are MIPs from laser scanning confocal microscopy. Scale bar = 25 μm. **(D)** Quantification of *C*. Integrated intensities from RAB5+ area are graphed for control and IFT20 KD HDLECs separately and combined. Intensities are graphed relative to average control HDLEC RAB5 intensity in the 45 min starve condition. Data points represent the average of three FOVs within one technical replicate. Each of three independent experiments included two or three technical replicates (three FOVs each) per experimental condition. Quantification is representative of 300+ control and 300+ IFT20 KD HDLECs. *p<0.05, **p<0.005, ***p<0.001.

### RAB5+ endosomes accumulate in IFT20 KD human dermal lymphatic endothelial cells

Members of the RAB family of small RAS-like GTPases regulate vesicular trafficking by localizing to the cytosolic leaflet of specific target membranes and recruiting effector proteins(Hutagalung and Novick, 2011; Naslavsky and Caplan, 2018). This steers cargo proteins through various vesicular compartments to their functional destination or for degradation. RAB5A localizes to the plasma membrane and early endosome and recruits effectors such as EEA1 (early endosome antigen 1) to early endosomes, where endocytosed proteins undergo sorting(Simonsen et al., 1999). IFT20 colocalizes with RAB5 in other cell types and is required for advancing cargo out of early endosomes such as recycling of membrane receptors and integrins(Finetti et al., 2014; Su et al., 2020). We first measured the basal level of RAB5A+ (hereafter referred to as RAB5) endosome formation in control and IFT20 KD HDLECs in homeostasis. HDLECs in homeostasis are cultured in a rich endothelial growth medium (EGM-2MV), supporting a moderate basal level of endocytic trafficking. As reported in other cell types, RAB5+ structures accumulated in IFT20 KDs compared to control cells, implying a defect in endosome maturation (**Figure 4A-B**). In control HDLECs expressing IFT20, we also noted punctate staining of IFT20 suggestive of localization to a vesicular compartment, similar to what we observed in mLECs (**Figure 4A** *top left panel*, **Figure 3A** *left panel*). In T cells, IFT20 KD did not inhibit internalization of the T cell receptor, transferrin receptor, or CXCR4, corroborating our data that IFT20 KD does not affect internalization of membrane proteins or formation of early endosomes(Finetti et al., 2014).

VEGF-C is the canonical growth factor that drives lymphangiogenesis, including in acute corneal inflammation. Above, we showed that corneal lymphangiogenesis was increased in LEC-specific IFT20 KO mice and that IFT20 KO mLECs migrated more strongly in response to VEGF-C as a chemoattractant in a transwell migration assay. This suggests a regulatory role for IFT20 in signaling downstream of VEGF-C. Here, we stimulated LECs with VEGF-C followed by VEGF-C washout and measured RAB5+ endosome formation to determine the impact of IFT20 on the dynamics of RAB5 endosome formation and resolution (**Figure 4C-D**). After 45 min, VEGF-C-stimulated IFT20 KD HDLECs accumulated RAB5+ endosomes similar to control HDLECs, suggesting that IFT20 does not regulate endocytosis or the recruitment of RAB5 to early endosomes. However, 90 min following washout of VEGF-C, the level of RAB5+ endosomes was strongly increased in IFT20 KD HDLECs, while this began to decrease toward baseline in control cells. These data suggest that IFT20 supports maturation of early endosomes, while endocytosis and early endosome formation are independent of IFT20 in HDLECs.

### VEGF-C stimulation increases IFT20/RAB5 colocalization in HDLECs

Previous studies in other cell types have shown that IFT20 localizes to the RAB5+ early endosome and supports cargo sorting to other vesicular compartments(Finetti et al., 2014, 2021; Su et al., 2020). Given the punctate IFT20 staining we observed and the effects of IFT20 KD on RAB5+ endosome dynamics, we hypothesized that IFT20 would localize to the RAB5+ compartment in HDLECs, particularly after VEGF-C stimulation. As in our previous experiments, control HDLECs in homeostasis contained at least two pools of IFT20 detectable by confocal immunofluorescence microscopy. We identified a perinuclear pool likely corresponding to the pool identified at the Golgi apparatus in other cell types(Follit et al., 2008; Galgano et al., 2017; Nishita et al., 2017; Finetti et al., 2021) as well as a significant pool made up of punctate structures distributed throughout the cytoplasm (**Figure 5A**). RAB5 immunostaining partially overlapped with this vesicular pool of IFT20, suggesting that IFT20 localizes to the early endosome in HDLECs during homeostasis. As expected, 24 h serum starvation of control HDLECs reduced the number of small RAB5+ endosomes, indicating a reduction in basal endocytosis, and the punctate pool of IFT20 was largely lost (**Figure 5B, C**). Perinuclear/Golgi-localized IFT20 and local large RAB5+ sorting endosomes appeared unchanged. RAB5 colocalized with approximately 15% of the total IFT20+ area after serum starvation. Stimulation of HDLECs with VEGF-C for 6 h resulted in the return of both small RAB5+ structures and punctate IFT20 staining throughout the cytoplasm. Colocalization between RAB5 and IFT20 was significantly increased, with approximately 40% of the total IFT20+ area colocalized with RAB5. By 1.5 h after VEGF-C washout and incubation of cells in basal media, the colocalization of RAB5 and IFT20 had significantly declined, and this was returned to baseline by 3 h. These data demonstrate that IFT20 is recruited to the early endosome by VEGF-C stimulation and suggest that it may regulate sorting of internalized proteins in HDLECs.

**Figure 5.**
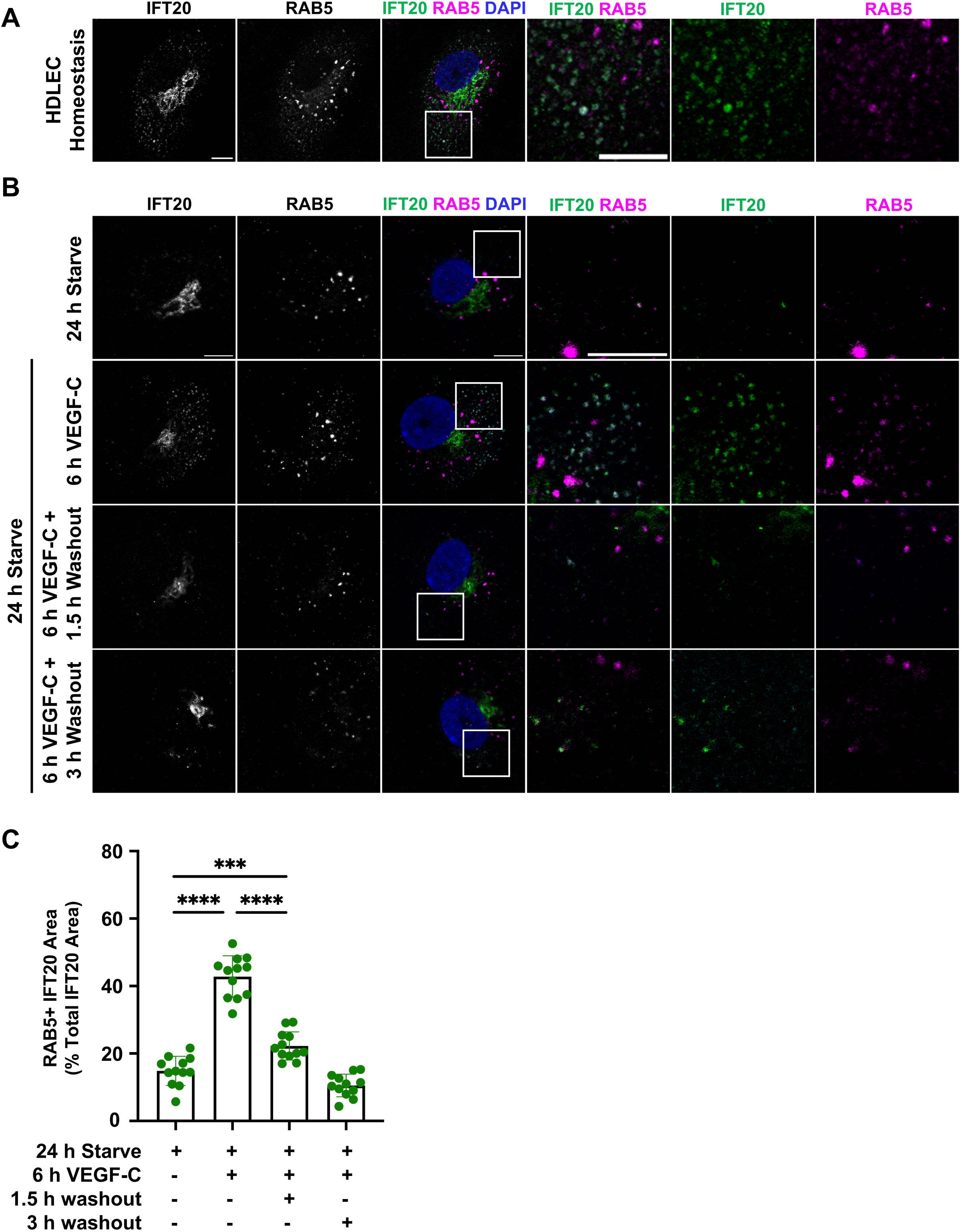
VEGF-C stimulation increases IFT20/RAB5 colocalization in HDLECs. **(A)** Non-serum starved control HDLECs were immunostained for IFT20 (*green*) and RAB5 (*magenta*). Scale bars = 10 μm. **(B)** Control HDLECs were serum starved for 24 h in 1/5 EGM-2MV and 4/5 EBM-2 (hereafter, starving media) and then treated for 6 h with 2 μg/mL VEGF-C in starving media. Cells were then fixed or placed into EBM-2 basal media for 1.5 or 3 h washout. IFT20 (*green*) and RAB5 (*magenta*) were detected by immunofluorescence microscopy. **(A-B)** Micrographs are single 0.3 μm z-slices from laser scanning confocal microscopy. DAPI (*blue*). Scale bars = 10 μm. **(C)** Quantification of *B*. RAB5+ IFT20+ area is graphed as a percentage of total IFT20+ area per FOV representing one of two biological replicates. Quantification is representative of 100+ control and 100+ IFT20 KD HDLECs. ***p<0.001, ****p<0.0001.

### IFT20 KD HDLECs accumulate VE-cadherin+ RAB5+ endosomes during VEGF-C stimulation and washout

Our *in vivo* and *in vitro* immunofluorescence experiments demonstrated that IFT20 is essential for proper inter-LEC junction organization. Based on the colocalization of IFT20 and RAB5 following VEGF-C stimulation, we hypothesized that IFT20 regulates junction architecture by controlling the vesicular trafficking of VE-cadherin following endocytosis. VEGF-C stimulation of VEGFR-3 has been shown to induce phosphorylation of the cytoplasmic tail of VE-cadherin, promoting its endocytosis and the breakdown of adherens junctions(Sung et al., 2022). Upon phosphorylation by Src, VE-cadherin undergoes clathrin-mediated endocytosis and then localizes to the RAB5+ early endosome(Wallez et al., 2007; Hebda et al., 2013; Yang et al., 2015). To determine how the absence of IFT20 affects cell-cell junction continuity, we stimulated control and IFT20 KD HDLECs with VEGF-C and immunostained for VE-cadherin (**Figure 6A**). After 24 h serum starve, IFT20 KD HDLECs displayed discontinuous VE-cadherin staining at cell-cell junctions, while this pattern was linear and continuous in control cells. As expected, after stimulation with VEGF-C for 6 h, breakdown of adherens junctions in both control and IFT20 KD LECs was observed, though this was more pronounced in IFT20 KD LECs. At 1.5 h post washout of VEGF-C, control cell junctions had begun to reestablish linearity and continuity, while those of IFT20 KD cells remained disorganized and discontinuous. Quantification of VE-cadherin granularity confirmed this result (**Figure 6B**). These results suggest that the trafficking function of IFT20 is important to reestablish interendothelial adherens junctions after VEGF-C-stimulated junctional breakdown.

**Figure 6.**
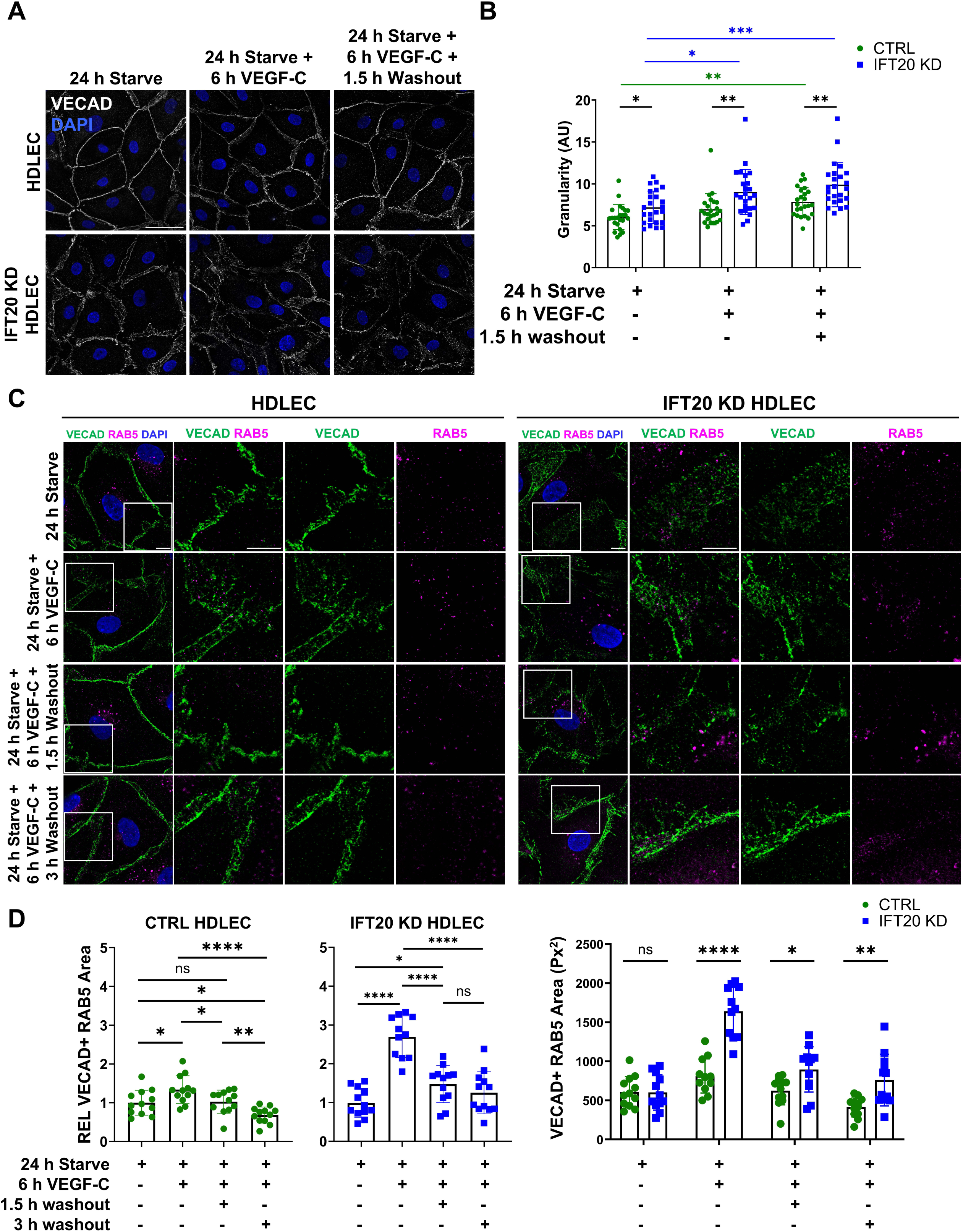
IFT20 KD HDLECs accumulate VE-cadherin+ RAB5+ endosomes during VEGF-C stimulation and washout. **(A)** Control and IFT20 KD HDLECs were serum starved for 24 h in in 1/5 EGM-2MV and 4/5 EBM-2 (hereafter, starving media). Cells were then treated with 2 μg/mL VEGF-C for 6 h in starving media and fixed or placed in EBM-2 basal media for 1.5 or 3 h washout. VE-cadherin (*white*) was detected by immunofluorescence microscopy. Scale bar = 50 μm. **(B)** Quantification of granularity to measure breakdown in VE-cadherin cell-cell junctions from *A*. Each data point represents the average granularity of a single FOV normalized per cell from each of two biological replicates. *p<0.05, **p<0.005. **(C)** Control and IFT20 KD HDLECs were serum starved for 24 h in starving media. Cells were then treated with 2 μg/mL VEGF-C for 6 h in starving media and fixed or placed in EBM-2 basal media for 1.5 or 3 h washout. VE-cadherin (*green*) and RAB5 (*magenta*) were detected by immunofluorescence microscopy. Scale bar = 10 μm. **(D)** Quantification of VE-cadherin+ RAB5+ area from *C*. Graphs show one representative biological replicate of two, each comprising two technical replicates with 100+ cells per condition. In graphs depicting individual CTRL or KD data, data is graphed relative to control 24 h starve treatment (column 1). In the *right graph* where the two cell lines are combined, data is graphed as total area (px^2^) for both datasets. *p<0.05, **p<0.005, ***p<0.001, ****p<0.0001. **(A, C)** Micrographs are single 0.3 μm z-slices from laser scanning confocal microscopy. DAPI (*blue*).

Our previous experiments demonstrated that IFT20 localizes to a RAB5+ compartment and that RAB5+ structures accumulate in IFT20 KD HDLECs after VEGF-C stimulation. Based on the disrupted VE-cadherin localization in IFT20 KD cells, we hypothesized that IFT20 regulates reestablishment of cell-cell junctions by promoting VE-cadherin traffic after VEGF-C-stimulated endocytosis. We serum starved control and IFT20 KD HDLECs, treated them with VEGF-C for 6 h, and then assayed colocalization of VE-cadherin and RAB5 1.5 h and 3 h after VEGF-C washout and incubation in basal media. Both control and IFT20 KD HDLECs displayed increased VE-cadherin+ RAB5+ area after 6 h treatment with VEGF-C, but this was much higher in IFT20 KD cells (**Figure 6C, D**). At 1.5 h after washout of VEGF-C, colocalization of VE-cadherin and RAB5 had returned to baseline in control cells, but this remained elevated in IFT20 KD cells at both 1.5 h and 3 h after VEGF-C washout. Thus, these results are consistent with a model in which VE-cadherin remains sequestered in a RAB5+ compartment for an extended period in the absence of IFT20 after removal of VEGF-C stimulation.

### IFT20 KD HDLECs exhibit increased and sustained VEGFR-3 signaling in response to VEGF-C treatment

Sustained VEGFR-2/3 signaling is thought to emanate from internalized receptors signaling from an endosomal compartment(Lampugnani et al., 2006; Simons, 2012; Sung et al., 2022). VEGFR-3 localizes to a RAB5C+ EEA1+ endosomal compartment in LECs after VEGF-C stimulation(Korhonen et al., 2022). VE-cadherin has been shown to directly associate with VEGFR-2/3, limiting receptor endocytosis and thereby restricting signaling(Coon et al., 2015). Phosphorylation of VE-cadherin stimulates its endocytosis and is thought to remove its repression of VEGFR-3 endocytosis and signaling(Sung et al., 2022). The extended latency of VE-cadherin in intracellular compartments in IFT20 KD cells led us to hypothesize that the increased lymphangiogenesis and migration phenotypes we observed *in vitro* and *in vivo* are due to increased VEGFR-3 signaling from endosomes. While we were unsuccessful in our attempts to visualize phosphorylated VEGFR-3 by immunofluorescence microscopy, we used quantitative western blotting to determine how control and IFT20 KD cells respond to VEGF-C stimulation and washout after serum starvation (**Figure 7**). The phospho-VEGFR-3 (Y1230) antibody we used recognized two forms at approximately 195 and 125 kDa(Pajusola et al., 1994; Bando et al., 2004; Breslin et al., 2007). Neither phosphorylated form was observed in serum-starved control or IFT20 KD cells. In control cells, VEGFR-3 phosphorylation peaked after 15 min VEGF-C stimulation and was reduced after 30 min stimulation and 30 min washout. Similarly, we observed strong increases in both forms with 15 min VEGF-C stimulation in IFT20 KD cells. However, in IFT20 KD cells, levels continued to rise through the 30 min time point. We observed similar results when analyzing phosphorylation of VE-cadherin at Y685, which is phosphorylated by Src(Wallez et al., 2007). IFT20 KD cells exhibited sustained VE-cadherin Y685 phosphorylation after 30 min of stimulation. Phosphorylation events on ERK and AKT occur downstream of VEGFR-3 activation and are important for HDLEC migration(Deng et al., 2015). Phospho-ERK and phospho-AKT levels showed similar trends as phospho-VEGFR-3 and phospho-VE-cadherin, with greater increases in IFT20 KD cells. These data suggest that Src-mediated phosphorylation of VE-cadherin enables extended VEGF-C/VEGFR-3 signaling in the absence of IFT20, likely from an endosomal compartment, and that this may promote LEC migration and lymphangiogenesis via ERK and/or AKT signaling.

**Figure 7.**
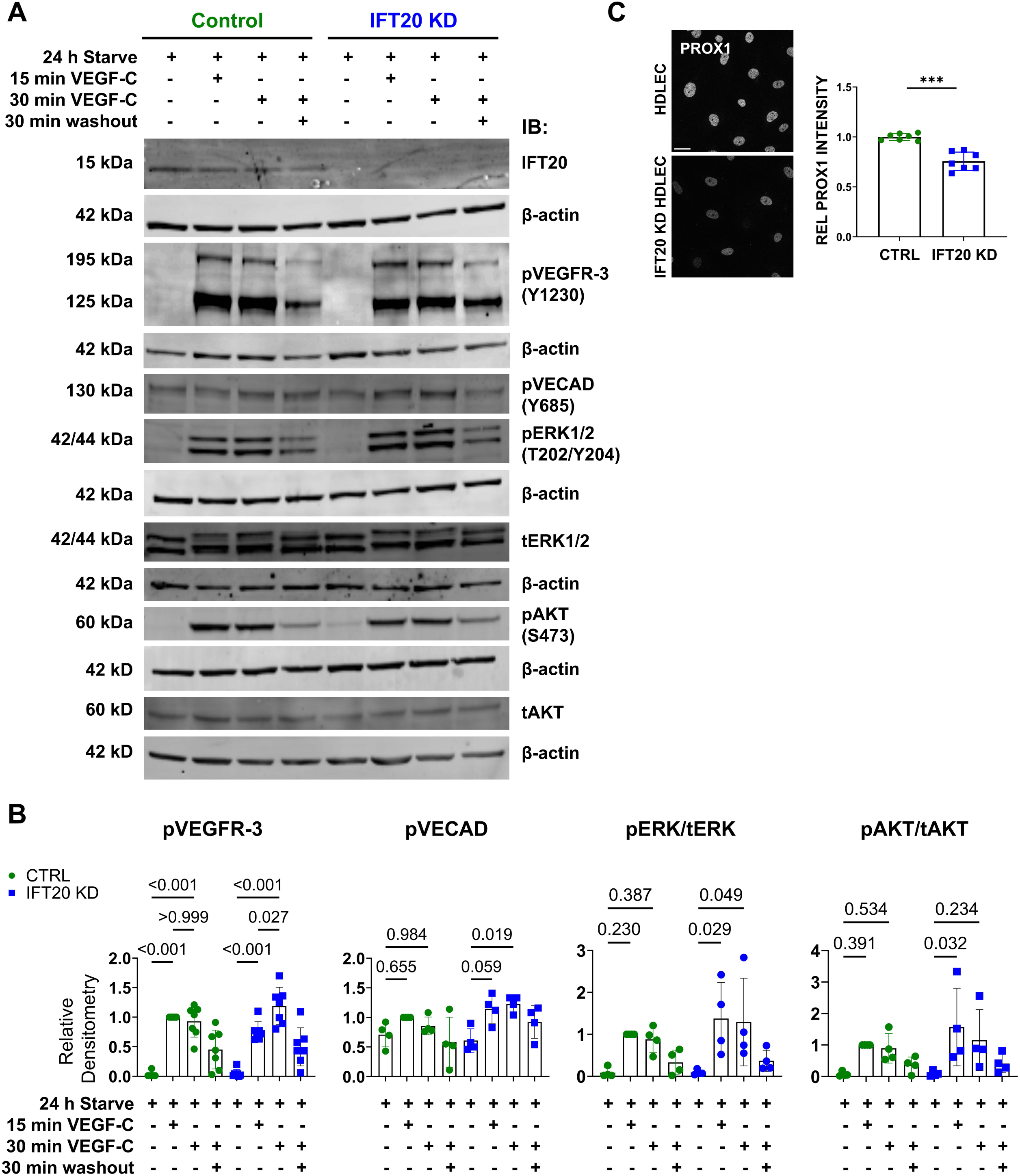
IFT20 KD HDLECs exhibit increased and sustained VEGFR-3 signaling in response to VEGF-C treatment. **(A)** Control and IFT20 KD HDLECs were serum starved for 24 h in 1/5 EGM-2MV and 4/5 EBM-2 (hereafter, starving media). Cells were then treated with 1 μg/mL VEGF-C for 15 or 30 min in starving media and lysed or placed in EBM-2 basal media for 30 min washout. Lysates were subjected to SDS-PAGE and western blot for the indicated proteins. β-actin is shown as a loading control below each band, respectively. β-actin shown in the 7^th^ row applies to both pVECAD and pERK1/2. **(B)** Protein levels were quantified by densitometry including background subtraction locally around each band and normalized to β-actin and, where indicated, levels of total protein (tERK1/2, tAKT). Each data point represents a single densitometry value from a total of four biological replicates. Data are graphed relative to column 2 (control cells stimulated with VEGF-C for 15 min) set equal to 1. **(C)** Control and IFT20 KD HDLECs in homeostasis were immunostained for PROX1. Scale bar = 25 μm. Quantification of integrated intensity from PROX1+ area in control and IFT20 KD HDLECs. Intensities are graphed relative to average control HDLEC PROX1 intensity. Data points represent the average from three FOVs in one technical replicate. Each of three independent experiments included two or three technical replicates (three FOVs each) per experimental condition. Quantification is representative of 300+ control and 300+ IFT20 KD HDLECs. *p<0.05, **p<0.005, ***p<0.001.

Through the course of our experiments, we also identified downregulation of PROX1 in IFT20 KD HDLECs under homeostasis growth conditions (**Figure 6C**). PROX1 is a transcription factor that regulates the expression of other genes required for LEC specification and fate, including those required for valve development (Sabine et al., 2012). Downregulation of PROX1 could result in a shift of these cells away from their terminally differentiated LEC phenotype, supporting loss of lymphatic vascular integrity and activation of non-quiescent or EMT characteristics in LECs. More studies are needed to understand how depletion of IFT20 results in downregulation of PROX1 protein and the consequences of this for LEC identity and function.

## Discussion

In this study, we assessed the mechanisms by which IFT20 regulates lymphatic vessel structure and function. We found that IFT20 controls vesicular trafficking of VE-cadherin. VE-cadherin is a key component of interendothelial adherens junctions and regulates VEGFR-3 signaling. IFT20 depletion caused accumulation of VE-cadherin in RAB5+ endosomes and concomitant breakdown of cell-cell contacts. This was correlated with increased LEC migration, increased inflammation-associated lymphangiogenesis, and increased lymphatic vessel leakiness. Localization of VE-cadherin in endosomes enhanced VEGFR-3 signaling, with an increase in phosphorylation of VEGFR-3, ERK1/2, and AKT. This study elucidates the function of an IFT protein in LECs and provides mechanistic insight into the processes that regulate lymphatic endothelial cell-cell junctions and lymphangiogenic signaling (**Figure 8**).

**Figure 8.**
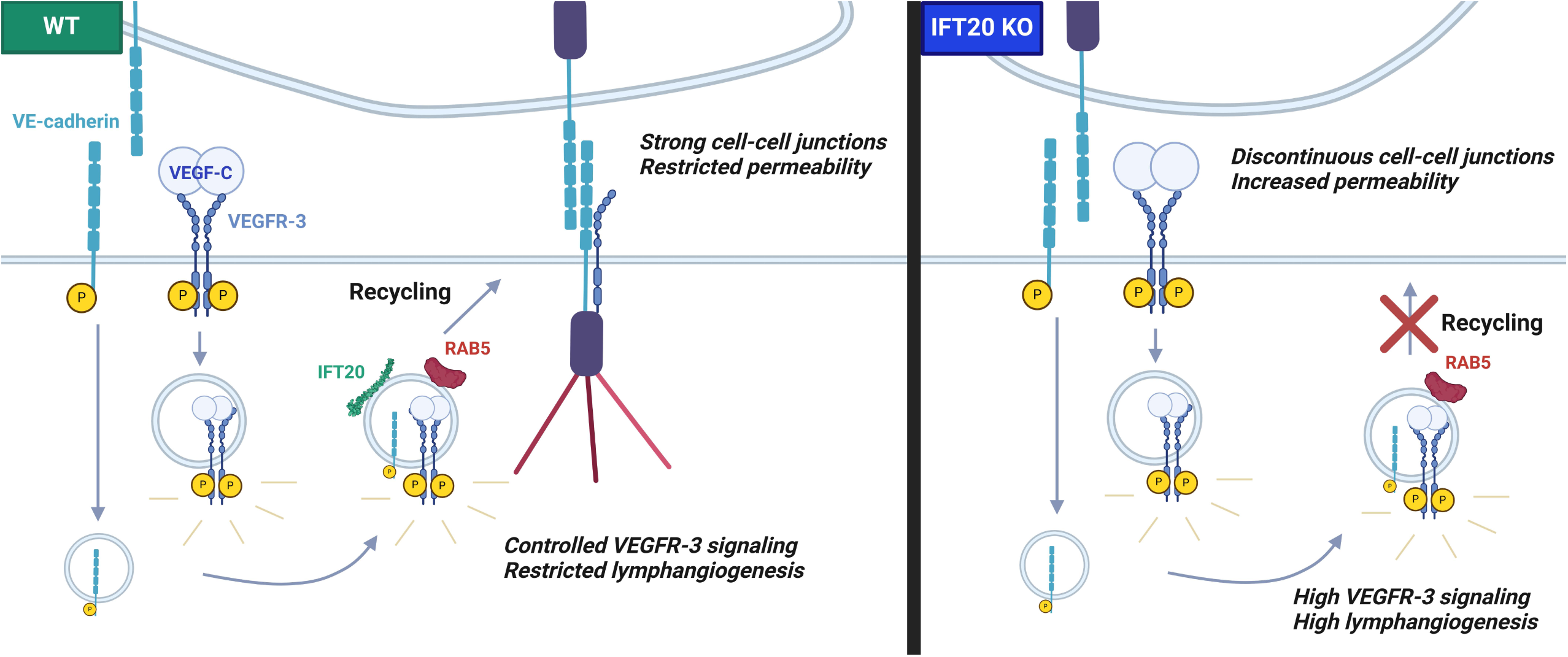
IFT20 controls VEGF-C signaling in LECs by regulating endocytic trafficking of VE-cadherin. Model: In the absence of IFT20, VE-cadherin accumulates in RAB5+ endosomes. This results in discontinuous cell-cell junctions and increased lymphatic vessel permeability. Sequestration of VE-cadherin inside the cell enables internalization and sustained signaling of VEGFR-3, promoting lymphangiogenesis. Model created with BioRender.com.

### IFT20 and Polarized Vesicular Traffic

IFT proteins are best characterized as members of complexes that transport cargo along the cytoskeletal axonemes of flagella, motile cilia, and nonmotile primary cilia. In addition to its intraciliary trafficking role as part of IFT complex B, IFT20 is unique among IFT proteins in its pattern of cellular localization and growing resume of extraciliary trafficking functions. Outside the primary cilium, IFT20 localizes to the base of the cilium, the Golgi apparatus, the centrosome, and vesicles(Follit et al., 2006). While the transport mechanisms that IFT20 participates in are nuanced for each cargo, a major theme is polarized recycling to specific plasma membrane regions. For example, IFT20 is required for traffic of ciliary membrane proteins such as polycystin-2 and rhodopsin to the primary cilium(Keady et al., 2011). IFT20 also regulates polarized traffic of planar cell polarity pathway protein Vangl2 to particular membrane locations(May-Simera et al., 2015). Furthermore, although nonciliated organisms do not express homologs of IFT20(Follit et al., 2006), mammalian cells that do not form primary cilia, such as T cells, do express IFT20, where it participates in trafficking pathways critical for cellular activation responses. In these cells, the vesicular-/Golgi-localized pool of IFT20 regulates cellular trafficking functions such as T cell receptor and integrin recycling(Finetti et al., 2009, 2014; Su et al., 2020). T cell receptor recycling to the immune synapse and integrin trafficking to the focal adhesion are endocytic trafficking pathways that rely on polarized IFT20-dependent vesicular transport out of RAB5+ endosomes.

Here, we propose that the interendothelial adherens junction is a similar structure. Our findings are consistent with a model in which VEGF-C stimulates VE-cadherin endocytosis and transit to the RAB5+ endosome, whereupon IFT20 promotes transit of VE-cadherin out of the RAB5+ endosome and polarized recycling back to cell-cell contacts. Our data suggest that IFT20 regulates VE-cadherin recycling to the adherens junction in an analogous manner to its regulation of T cell receptor recycling to the immune synapse. In both cases, IFT20 localizes to RAB5+ endosomes, and depletion of IFT20 causes accumulation of cargo in RAB5+ endosomes, suggesting that transit to RAB4+ or RAB11+ endosomes for recycling may be impaired. A recent study in human lung microvascular endothelial cells demonstrated that RAB11a is required for recycling of VE-cadherin(Yan et al., 2016). IFT20 is implicated in RAB11-dependent recycling of the T cell receptor, in concert with RAB8(Finetti et al., 2015). Further studies are needed to determine how IFT20 regulates trafficking of VE-cadherin beyond the RAB5+ endosome. IFT20 is required not only for recycling of the T cell receptor but also for delivery of the downstream signaling mediator LAT to the immune synapse(Finetti et al., 2014; Vivar et al., 2016). Future studies are needed to determine if IFT20 or other IFT proteins participate in non-vesicular trafficking of adaptor proteins such as catenins to adherens or other intercellular junctions.

Localization of IFT20 to the Golgi apparatus requires golgin GMAP210, where the two work in concert along with GM130 and VPS15 to regulate post-Golgi traffic (Follit et al., 2008; Stoetzel et al., 2016) and, in some cell types, Golgi polarization during migration(Aoki et al., 2019). However, unlike IFT20 KO embryos, GMAP210 KO embryos do not display edema(Follit et al., 2008). This suggests that localization and function of IFT20 at the Golgi apparatus is not critical for preventing lymphatic vessel dysfunction. Rather, it implicates a distinct trafficking role, such as recycling of cell surface proteins, in fluid homeostasis defects and edema. To our knowledge, lymphatic vessel development and remodeling in the absence of GMAP210 has not been studied and would be an interesting research avenue.

During T cell receptor recycling, IFT20 facilitates polarized exocytosis from recycling endosomes at the immune synapse (in complex with IFT88 and IFT57)(Finetti et al., 2009). A recent BioID proteomics study in HDLECs identified components of the exocyst complex as binding partners of VE-cadherin during adrenomedullin-stimulated junctional remodeling(Serafin et al., 2024). Previous work has shown that IFT20 directly interacts with Exo70 (human exocyst complex component 7 (EXOC7)) in the exocyst complex to regulate trafficking of ciliary proteins fibrocystin and polycystin-2(Monis et al., 2017). IFT20 also directly interacts with Sec10 (human EXOC5)(Fogelgren et al., 2011). Importantly, the junctional phenotype observed in EXOC5 KD HDLECs(Serafin et al., 2024) was highly reminiscent of the junctional phenotype we observed in IFT20 KD HDLECs. Further studies are needed to determine whether IFT20 participates in the predicted exocyst-mediated recycling of VE-cadherin.

In addition to exocyst components, it has been demonstrated that IFT20 functions with other proteins in various complexes, including but not restricted to other IFT proteins, in both ciliary and nonciliary transport pathways. For example, IFT20 directly interacts with IFT88, 57, 52(Follit et al., 2006), and IFT52 and 57 were implicated in T cell receptor trafficking along with IFT20(Finetti et al., 2014). A growing body of work also indicates that IFT20 interacts with other proteins to regulate cellular degradation pathways. IFT20 interacts with ATG16L1 to regulate cellular autophagy(Boukhalfa et al., 2021; Finetti et al., 2021) and regulates lysosome maturation through delivery of acid hydrolase enzymes via classical trafficking mechanisms such as retrograde transport of cation-independent mannose-6-phosphoate receptor and association with motor protein dynein (Pampliega et al., 2013; Zhang et al., 2016; Finetti et al., 2020, 2021). Future studies in LECs are necessary to understand the nuances of the IFT20 interactome and its role in ciliary vs. nonciliary processes.

### Implications for Lymphatic Valve Formation

The lymphatic system develops valves to promote unidirectional transport of lymph. This process depends on cellular polarization and migration events as LECs orient perpendicularly to the direction of lymph flow and form valve leaflets supported by ECM (Bazigou and Makinen, 2013; Yang et al., 2019). Lymphatic valvulogenesis requires proper adherens junction formation(Tatin et al., 2013), and valves contain high levels of VE-cadherin organized continuously in zipper-like junctions(Wang et al., 2016). Deletion of VE-cadherin(Hägerling et al., 2018; Yang et al., 2019) or connexin-43(Kanady et al., 2011) can prevent valve formation or cause valve regression. The retrograde lymph flow that we observed in our ear skin model suggests a possible valve defect (**Figure 2E**). Similarly, we previously reported reflux of red blood cells into lymphatic vessels in IFT20 KO embryos(Paulson et al., 2021), suggesting a defect in lymphovenous valve integrity(Yang and Oliver, 2014). We also observed downregulation of PROX1 in IFT20 KD HDLECs (**Figure 7C**). PROX1 upregulation is a key step in the establishment of a valve territory(Cha et al., 2016), and its downregulation here may indicate an additional IFT20- or primary cilia-based mode of valve regulation. Future studies should directly assess the effects of IFT20 depletion or primary cilia assembly inhibition on lymphatic valve formation, maintenance, and function.

### VE-cadherin and VEGFR-3 Signaling

VEGF family protein ligands bind VEGFRs and regulate many facets of lymphatic and blood endothelial cell biology. Lymphangiogenesis can be stimulated by VEGF-C binding VEGFR-3 or VEGFR-2, which stimulates LEC proliferation and migration(Joukov et al., 1996; Mäkinen et al., 2001; Simons et al., 2016). VEGFR-3 signaling is required for the establishment of VE-cadherin button junctions as lymphatic collecting vessels mature but is not required for their maintenance(Jannaway et al., 2023). A recent study elegantly demonstrated the reciprocal inhibition of VE-cadherin and VEGFR-3 by their sequestration the cell surface (Sung et al., 2022). During endothelial cell quiescence in mature blood and lymphatic vessels, VE-cadherin molecules on adjacent cells provide strong homotypic adhesion that inhibits cell sprouting and migration(Bentley et al., 2009), while loss of VE-cadherin caused fragmentation of vasculature(Yang et al., 2019). Membrane-localized VE-cadherin directly promotes the maintenance of VEGFR-2 or VEGFR-3 at the membrane, thereby limiting their signaling and promoting vascular plexus homeostasis(Calera et al., 2004; Sung et al., 2022). Endocytosis of VE-cadherin to EEA1+ endosomes is stimulated when VEGF-C/VEGFR-2/3 signaling activates Src kinase to phosphorylate Y685 on the VE-cadherin cytoplasmic domain(Wallez et al., 2007; Sung et al., 2022). Internalization of VE-cadherin relieves the repression of VEGFR-2/3 endocytosis, enabling these receptors to robustly signal from the RAB5C+ EEA1+ endosome after VEGF-C stimulation(Lampugnani et al., 2006; Simons, 2012; Korhonen et al., 2022). Expression of an α-catenin:VE-cadherin fusion protein resulted in stabilization of VE-cadherin at the cell surface(Sung et al., 2022). Mice expressing this construct developed embryonic edema due to increased VEGFR-3 residence at the cell surface, which restricted optimal signaling from the endosome and stunted lymphatic development. Conversely, haploinsufficiency of VE-cadherin rescued the edema defect in VEGFR-3 KO mice. This suggests that VE-cadherin inhibition of VEGFR-3 signaling by sequestering it at the membrane is analagous to limiting its signaling by depleting VEGFR-3 directly. VE-cadherin KO lymphatics displayed high levels of pVEGFR-3 and were hyperplastic in mesentery, a tissue with strong VEGF-C expression at the time point of analysis, indicating that these VE-cadherin KO vessels were hypersensitive to VEGF-C stimulation, rather than insensitive(Hägerling et al., 2018). Our data agree with and extend these studies, showing that deletion of IFT20 promotes VE-cadherin button junction formation (**Figures 1, 3, 6**) and lymph leakage (**Figure 2**). VEGF-C stimulation enhances these effects, both *in vivo* during corneal inflammation and *in vitro*, and correlates with enhanced phosphorylation of VEGFR-3, downstream activation of ERK and AKT (**Figure 7**), and increased LEC migration (**Figure 3**) and lymphangiogenesis (**Figure 1**).

## Acknowledgements

We thank Mason Crow, Caden Johnson, Hannah Polejewski, Taylor Schallenkamp, and Nora Smestad for their help with experiments. We gratefully acknowledge Dr. J. Steven Alexander (Louisiana State University Health Shreveport) for the SV-LEC cell line, Dr. Gregory J. Pazour (University of Massachusetts Chan Medical School) for the *Ift20*^fl/fl^ mice, and Dr. Pazour and Dr. Rachel A. Willand-Charnley (South Dakota State University) for the IFT20 CRISPR reagents. This material is based upon work conducted using the South Dakota State University Functional Genomics Core Facility (RRID:SCR_023786) supported in part by the National Science Foundation/EPSCoR Grant No. 0091948, the South Dakota Agricultural Experiment Station, and by the State of South Dakota. Research reported in this publication was supported by the National Institute of General Medical Sciences of the National Institutes of Health under Award Numbers P20GM135008 and R15GM140458. The content is solely the responsibility of the authors and does not necessarily represent the official views of the National Institutes of Health.

## Author Contributions

Conceptualization: AM, DP, DMF

Methodology: AM, DP, DMF

Formal analysis: AM, DP, DMF

Investigation: AM, DP, ZL, LK, JP, SL, DMF

Resources: DMF

Writing – Original Draft: AM, DP, DMF

Writing – Review and Editing: DP, AM, ZL, LK, JP, SL, DMF

Supervision: DMF

Project administration: DMF

Funding acquisition: DMF

## Methods

### Experimental Model and Study Participant Details

#### Mice

All mouse work was approved by the Institutional Animal Care and Use Committee of South Dakota State University and carried out in accordance with the Association for Research in Vision and Ophthalmology Statement for the Use of Animals in Ophthalmic and Vision Research. C57BL/6J mice were obtained from Jackson Laboratories (strain #000664). LYVE1Cre mice were obtained from Jackson Laboratories (strain #012601)(Pham et al., 2010) and maintained as homozygotes. IFT20^fl/fl^ mice were a kind gift from Gregory J. Pazour (University of Massachusetts Chan Medical School) and maintained as homozygotes. IFT20^fl/fl^ mice were crossed to LYVE1Cre mice to generate an intermediate generation bearing one floxed copy of *Ift20* and positive for Cre recombinase. This intermediate generation was crossed to IFT20^fl/fl^ mice to generate experimental mice with two floxed copies of *Ift20* and positive for Cre recombinase. Littermates bearing one floxed copy of *Ift20* and negative for Cre were used as controls. LYVE1Cre is active in LECs and has some leaky expression also in blood endothelial cells, as previously observed in our laboratory when crossing LYVE1Cre mice to a tdTomato reporter line (Jackson Laboratories #007914). Mice were housed in groups whenever possible with standard husbandry. Mice were used between 6 and 35 weeks of age. Both sexes were used and data were not analyzed separately by sex due to the extensive breeding required to generate experimental mice.

#### Cell Lines

##### Development of mLEC IFT20 KO Cell Line

Immortalized mouse mesenteric lymphatic endothelial cells (SV-LEC or mLEC) were a kind gift from J. Steven Alexander (sex not reported) (Ando et al., 2005). mLECs were routinely cultured with Dulbecco’s Modified Eagle’s Medium (DMEM) (Fisher Scientific, 11-995-065) supplemented with 10% fetal bovine serum (Fisher Scientific, 10-082-147) and 1X penicillin/streptomycin (Fisher Scientific, 15-140-122) at 37°C and 5% CO_2_. CRISPR-Cas9 was used to knockout *Ift20* from mLECs using lentiviral vector transduction and clonal selection. Guide sequence: GACTGAACAAGCTCCGAGTGT. 1 μL of IFT20 Cas9 Lentiviral vector PD205.1 plasmid (a kind gift from Dr. Gregory J. Pazour, University of Massachusetts Chan Medical School) was transformed into TOP10 *E. coli* competent cells (ThermoFisher, C404010), incubated at 42°C for 45 s, and then transferred to ice for 3-5 min. Transformed cells were grown in 100 μg/mL ampicillin-supplemented Luria Bertani media. The next day cells were lysed, and the plasmid was extracted following the QIAGEN Miniprep kit protocol. 293T packaging cells were then transfected with 1.5 mg PD205.1, 0.1 mg of GAG/POL, REV and TAT packaging plasmids (gifts from Dr. Rachel Willand-Charnley, South Dakota State University) suspended with Mirus-293T lipid in Opti-MEM. The plasmid solution was added dropwise to 70-80% confluent 293T packaging cells. Media was changed after 24 h. After 72 h the supernatant was collected, aliquoted, and frozen. The virus-containing supernatant was thawed, spun at 4,900 RPM at room temperature, and then added to the mLEC cells at approximately 50% confluency. After 24 h, the supernatant was discarded and fresh DMEM media containing 1.5 μg/mL of puromycin was added to the cells. The cells were kept under puromycin selection until non-transduced control mLECs were dead. To obtain a pure population of KO cells, we performed clonal selection. 50 cells were seeded on a 10 cm plate containing 10 mL media. Individual colonies were selected, grown, and analyzed for IFT20 expression. Knockout was confirmed via immunofluorescence staining. Parental cells were used as controls in all assays.

##### siRNA Knockdown of IFT20 in HDLECs

Adult human dermal lymphatic endothelial cells (HDLECs) from healthy donors were purchased from PromoCell (C-12217). HDLECs from PromoCell are CD31+ and Podoplanin+. HDLECs were routinely cultured in Endothelial Cell Growth Media MV2 (EGM-MV2) (PromoCell, C-39226) at 37°C and 5% CO_2_. ON-TARGETplus Human IFT20 siRNA Pool (L-017747-01-0005) and ON-TARGETplus Non-targeting Control Pool (D-001810-10-05) were purchased from Horizon Discovery Dharmacon. The IFT20 siRNA pool contained siRNAs corresponding with the following six accession references: NM_001267774, NM_001267775, NM_001267776, NM_001267777, NM_001267778, and NM_174887. An IFT20 siRNA knockdown (KD) protocol for HDLECs was developed from Invitrogen’s siRNA into HUVEC Cells Using Lipofectamine™ RNAiMAX protocol. Briefly, HDLECs were cultured in 25 cm^2^ tissue culture-treated flasks to 70-80% confluency. 65 pmol of IFT20 siRNA or non-targeting control siRNA along with 19.5 μL Lipofectamine™ RNAiMAX (Fisher Scientific, 13778075) transfection reagent was diluted in 1.3 mL Opti-MEM® I Reduced Serum Medium (Fisher Scientific, 31985062). The siRNA-Lipofectamine dilution was incubated at room temperature for 20 min and then added to the flask containing HDLECs and 6.5 mL antibiotic-free EGM-MV2. The final siRNA concentration used was 8.2 nM. The siRNA-Lipofectamine mixture incubated with HDLECs at least 12 h at 37°C and 5% CO_2_. IFT20 KD and control HDLECs were then utilized in various functional and biochemical assays within 72 h of siRNA transfection. Knockdown of IFT20 was confirmed via immunofluorescence staining in parallel with each assay and by western blot as shown in Figure 7. Cells treated with scrambled siRNA were used as controls in all assays.

## Method Details

### Inflammation-Induced Corneal Lymphangiogenesis

We used a model of acute corneal inflammation as previously described(Kelley et al., 2011, 2013; Fink et al., 2014). Mice were placed under general (ketamine 10 mg/kg, xylazine 1 mg/kg) and topical (proparacaine hydrochloride, USP, 0.5%) anesthesia. Four 10-0 proline monofilament sutures were placed in the cornea of one eye of healthy mice. Mice were provided with supportive care following surgery including subcutaneous administration of saline, topical treatment with antibiotic ophthalmic ointment, and recovery on a heating pad. After 12 days, mice were euthanized, and corneas were harvested for whole-mount immunofluorescence staining.

### Lymphatic Function Assay and Intravital Imaging

Mice were anesthetized with a mixture of ketamine (100 mg/kg) and xylazine (10 mg/kg). 0.5 or 4 μL TRITC-conjugated dextran (40 kDa) was injected into the ear skin using a pulled glass needle with approximate diameter 75 μm. Intravital imaging was performed using a Zeiss SteREO Discovery.V8 microscope with a DsRed filter and an AxioCam 503 high resolution digital monochrome camera.

### Whole-Mount Immunofluorescence Staining

Ears and corneas from adult mice were harvested for whole-mount staining after euthanasia. All incubations were performed at room temperature. Dorsal and ventral halves of ears were separated and fixed in 2% paraformaldehyde (PFA) for 2 h. Whole eyes were harvested and fixed in 1% PFA. Corneas were then dissected out of eyes and fixed for an additional 1 h in fresh 1% PFA. Ears and corneas were then incubated for at least 1 h in 1X PBS. Next, tissues were blocked and permeabilized for 1 h (corneas) or overnight (ears) in PBS++, containing 5.2% bovine serum albumin (BSA), 0.3% Triton X-100, and 0.2% sodium azide in 1X PBS, pH 7.4. Primary and secondary antibodies were diluted in PBS++ and incubated overnight. Following both primary and secondary incubations, three 1 h washes were completed in PBS+, containing 0.2% BSA, 0.3% Triton X-100, and 0.2% sodium azide in 1X PBS, pH 7.4. Samples were mounted in Fluoromount-G without DAPI (Fisher Scientific, OB100-01).

### mLEC and HDLEC Immunofluorescence

mLECs and HDLECs were cultured on uncoated glass coverslips or ibiTreat μ-slides IV 0.4 (ibidi, 80606). LECs were fixed with ice cold methanol (ARL13B IF) or 4% paraformaldehyde in PBS, pH 7.2 for 15 min. Following fixation, cells were stored in 1X PBS at 4°C until immunostained. LECs were permeabilized with 0.15% Triton X-100 in 1X PBS for 15 min at room temperature. Next, LECs were blocked using 1% BSA in 1X PBS for 30 min at room temperature. Primary antibodies were then diluted in 1% BSA-PBS and incubated for 1-2 h at room temperature under gentle rocking. Three 5-min washes with 1X PBS followed primary antibody incubation. Next, secondary antibodies were diluted 1:10,000 in 1% BSA-PBS and incubated for 45 min at room temperature under gentle rocking. Three 5-min washes with 1X PBS followed secondary antibody incubation. Finally, coverslips were mounted in Fluoromount G with DAPI (Fisher Scientific, OB010020). Ibidi μ-slides were kept hydrated with 1X PBS containing NucBlue (ThermoFisher, R37606). Cells were imaged via epifluorescence or laser scanning confocal microscopy.

### Transwell Migration Assay

Transwell migration stimulated with 0.2% FBS-DMEM (negative control), 10% FBS-DMEM (positive control), or 500 ng/mL VEGF-C (R&D systems, 9199-VC) in 0.2% FBS-DMEM was assessed in IFT20 KO and control mLECs. 40,000 cells suspended in 100 µL 0.2% FBS-DMEM were seeded in the upper chamber of transwell permeable supports within a 24-well plate (Corning, CLS3422) onto a pre-soaked, equilibrated polycarbonate, tissue culture-treated membrane with 8 μm pores. 500 µL of stimulus-containing media was placed in the lower chamber. After 24 h incubation at 37°C and 5% CO_2_, non-migratory cells were removed from the top of the membrane using a cotton swab. The membrane was then rinsed once with 1X PBS and fixed with methanol. After drying completely, membranes were mounted in Fluoromount G with DAPI and imaged via epifluorescence microscopy. DAPI-labeled nuclei of migratory cells were quantified using CellProfiler’s Identify Primary Objects function, and statistical analysis was completed using GraphPad Prism. Quantifications represent one of three biological replicates, where each data point represents one quantified membrane. Each biological replicate consisted of three technical replicates (three membranes) per stimulus for each cell line. Migration is graphed relative to 10% FBS-DMEM-stimulated control mLEC migration.

### VEGF-C Stimulation for Immunofluorescence and Western Blot

For immunofluorescence experiments, HDLECs were serum starved for 24 h in media containing 1/5 EGM-MV2 growth media and 4/5 EBM-2 basal media, referred to hereafter as starving media (PromoCell). Following serum starvation, HDLECs were stimulated with 2 μg/mL VEGF-C (R&D systems, 9199-VC) in starving media or fresh starving media alone (negative control) for 45 min at 37°C and 5% CO_2_. Following stimulation, wells were rinsed to washout VEGF-C and then cells were incubated in basal EBM-2 media for 90 min at 37°C and 5% CO_2_. At desired stimulation or recovery timepoint, HDLECs were fixed with 4% PFA in PBS, pH 7.2, and immunostained as described above. This protocol was then modified to study junctional remodeling over several hours. Briefly, HDLECs were serum starved for 24 h then stimulated with VEGF-C for 6 h. Following stimulation, VEGF-C was washed out and HDLECs were allowed to recover for 1.5 or 3 h before fixing and staining. For Western blot experiments, the same protocol was followed as above with the following exceptions. HDLECs were stimulated with 1 μg/mL VEGF-C (R&D systems, 9199-VC) in starving media. VEGF-C stimulation time points were 15 and 30 min. Washout time point was 30 min after 30 min stimulation.

Primary antibodies utilized in cell and tissue immunofluorescence staining include: rabbit anti-mouse LYVE-1 1:200 (abcam, ab33682), rat anti-mouse LYVE-1 1:50 (Santa Cruz, sc-65647), goat anti-human PROX-1 1:200 (R&D Systems, AF2727), rabbit anti-mouse PROX-1 1:500 (abcam, ab101851), mouse anti-mouse ARL13B 1:200 and 1:4,000 (Neuromab, 75-287), rabbit anti-mouse IFT20 1:1,000 (Proteintech, 13615-1-AP), rabbit anti-mouse ZO-1 1:1,000 (Proteintech, 21773-1-AP), goat anti-mouse VE-cadherin 1:200 (R&D systems, AF1002), rat anti-mouse VE-cadherin 1:1,000 (BD Bioscience, 555289), and rat anti-PECAM-1 1:1,000 (BD Pharmingen, 553370). Directly conjugated primary antibodies used in whole-mount tissue immunofluorescence include: rabbit anti-Rab5A-AF555 1:500 (abcam, ab311900). Secondary antibodies used in cell and tissue immunofluorescence at concentrations of 1:10,000 and 1:500, respectively, include: goat anti-mouse 488, donkey anti-rabbit 488, goat anti-mouse 555, chicken anti-goat 647, donkey anti-rabbit 555, donkey anti-rat 594, chicken anti-goat 488, goat anti-rat 550, and goat anti-rat 647. F-actin was labeled using phalloidin-AF647 (Invitrogen, Cat. No. A22287).

### Microscopy

A Zeiss Stemi 305 compact stereo dissecting microscope with a 1080P HP digital video camera was used to perform surgery and dissections. A Zeiss SteREO Discovery.V8 fluorescence stereomicroscope and Axiocam 503 monochrome camera were used to image mice and mouse tissues and administer dextran injections. Epifluorescence microscopy was performed using a DMI 4000 B microscope (Leica) with a KUBler CODIX EL6000 light source (Leica) and a Zyla 4.2 sCMOS camera (ANDOR). Epifluorescence microscopy was also performed utilizing a Nikon E800 upright microscope paired with a Zeiss Axiocam 503 monochrome camera. Laser scanning confocal microscopy was performed using a FLUOVIEW FV1200 scanning confocal microscope (Olympus) interfaced with an IX81 microscope (Olympus) equipped with 488/559/635 laser lines or a Leica LiaChroic Stellaris 5 confocal system equipped with 405/488/514/559/638 laser lines and 3 HyD spectral detectors. Z-slice thickness from confocal microscopy is indicated in figure legends. Microscopy was performed at room temperature.

### Western Blot

IFT20 KD or control confluent HDLECs from two 9.6 cm^2^ wells were lysed in 100 μL of lysis buffer consisting of RIPA buffer (Thermo Fisher Scientific, 89900), 0.5 M 1X EDTA (Thermo Fisher Scientific, 1861274), and 1X protease and phosphatase inhibitor cocktail (Thermo Fisher Scientific, 1861281). Lysates were cleared by centrifugation at 16,000 X g at 4°C for 20 min. To measure the concentration of protein in the cell lysates, the BCA protein assay kit (Thermo Fisher Scientific, 23225) was used. The absorbance of the BCA reagent-treated lysate at 562 nm was measured using an Epoch microplate spectrophotometer (Agilent BioTek Instruments) and analyzed using Gene 6 software (version 1.04). Lysis buffer was added to lysates to bring the volume of each sample up to 20 μL while keeping the concentration of all the samples the same. 7.5 μL of NuPAGE 4X sample buffer (Invitrogen, NP0007) and 3 μL of NuPAGE 10X sample reducing agent (Invitrogen, NP0009) were added to each 20 μL sample before boiling for 5 min. For separation of the proteins based on molecular weight, the samples were run on a NuPAGE 4-12% Bis-Tris gel (Invitrogen, NP0321BOX) followed by transfer to an Immobilon-FL PVDF membrane (Millipore Sigma, IPFL07810) using the XCell II^TM^ Blot Module (Invitrogen, EI9051). After transfer, membranes were probed with primary antibodies against phospho-VE-cadherin, phospho-VEGFR-3, phospho-AKT, phospho-ERK1/2, total AKT, total ERK1/2, IFT20 and loading control β-actin overnight at 4°C followed by Licor IRDye secondary antibodies for 1 h at room temperature. The membranes were imaged using a LI-COR Odyssey CLx Imager with near-infrared western blot detection and analyzed using the software LI-COR Image Studio (version 5.2).

The following antibodies were used for Western blot analysis at the indicated concentrations: Rb anti-IFT20 (Proteintech,13615-1-AP) 1:500, Ms anti-β-actin (Santa Cruz Biotechnology, sc-47778) 1:500, Rb anti-phospho-VEGFR-3 (120232, Covalab) 1:1,000, Rb anti-phospho-AKT (Cell Signaling Technology, 4360T) 1:1,000, Rb anti-phospho-VE-cadherin (EMD Millipore Corporation, ABT1760) 1:1,000, Rb anti-AKT (Cell Signaling Technology, 4685T) 1:1,000, Rb anti-ERK1/2 (Cell Signaling Technology, 4695T) 1:2,000, Rb anti-phospho-ERK1/2 (Cell Signaling Technology, 4370T) 1:1,000. Secondary antibodies include: donkey anti-mouse IRDye 800CW 1:15,000 (Licor, 926-32212) and donkey anti-rabbit IRDye 680RD 1:15,000 (Licor, 926-68073).

### Quantification and Statistical Analysis

Quantification methods are described in figure legends and results sections. Image processing, analysis, and quantifications were carried out in FIJI, CellProfiler, ZEN, or LI-COR Image Studio software. Data were analyzed using GraphPad Prism software using Student’s or Welch’s T-Test or one-way ANOVA with Tukey’s multiple comparisons test. Replicates are indicated in figure legends. Data is centered at the mean and error bars show standard deviation.

### Key Resources Table

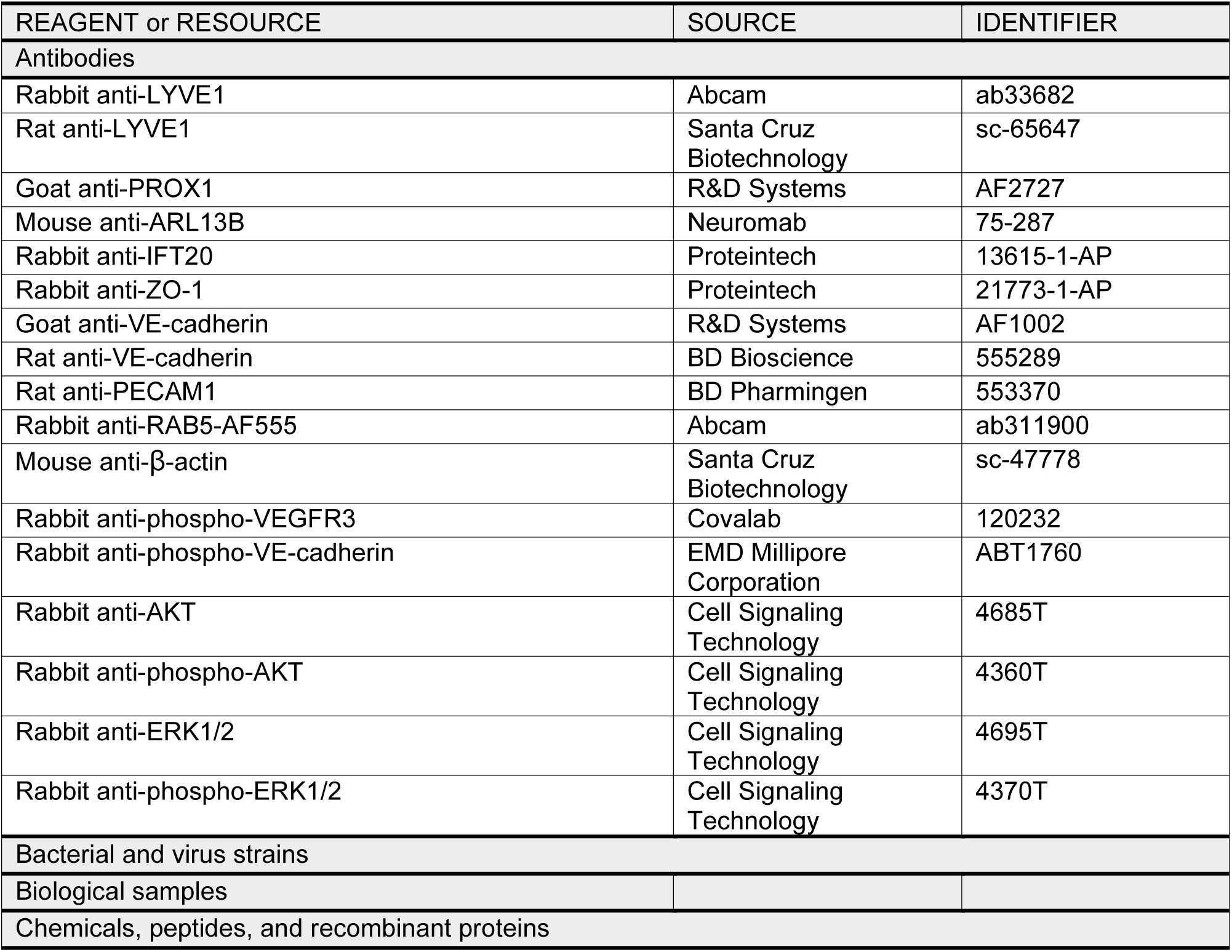
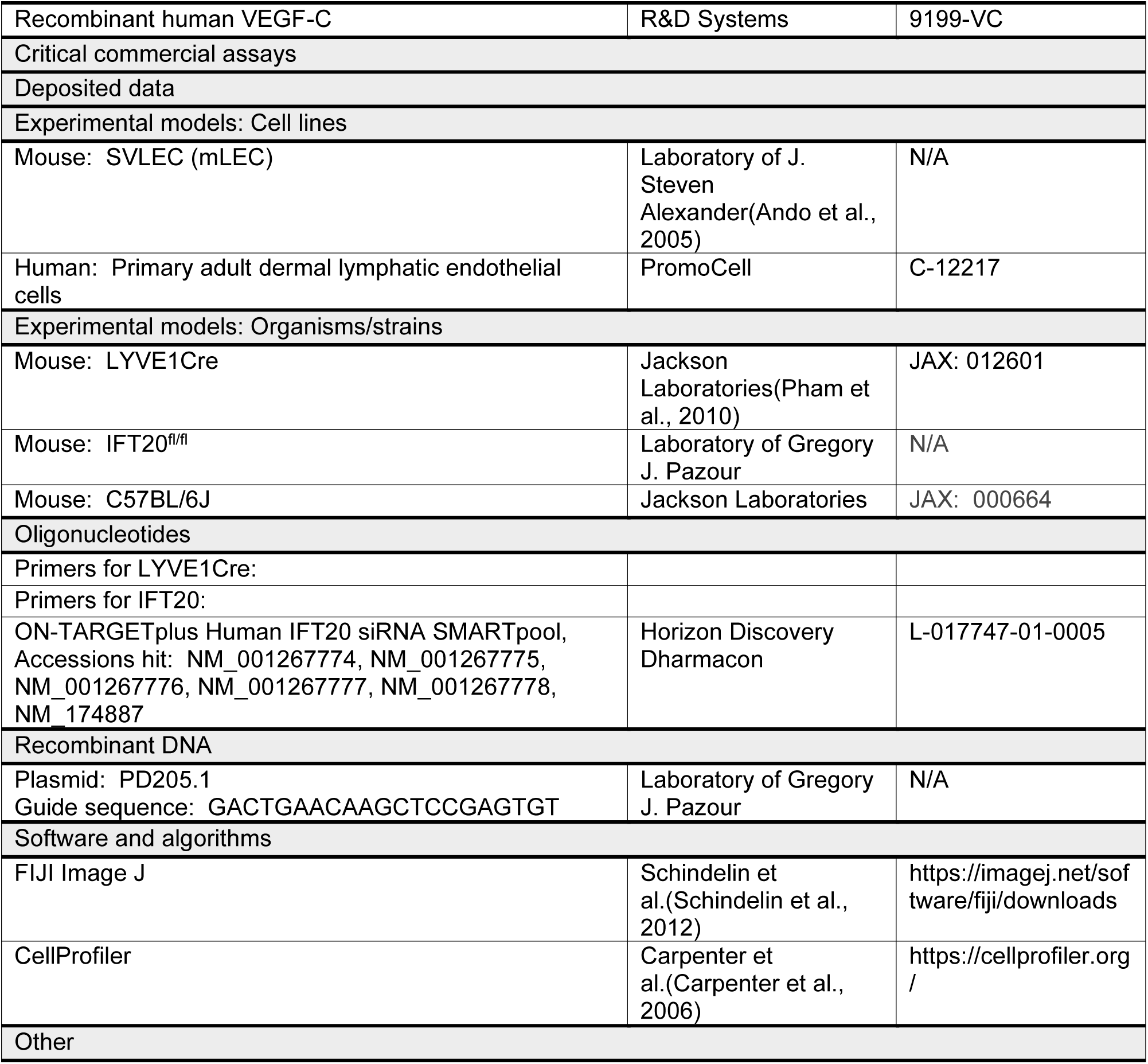

